# Function and regulation of an aldehyde dehydrogenase essential for ethanol and methanol metabolism of the yeast, *Komagataella phaffii*

**DOI:** 10.1101/2020.06.14.151365

**Authors:** Kamisetty Krishna Rao, Umakant Sahu, Pundi N Rangarajan

**Affiliations:** Department of Biochemistry, Indian Institute of Science, Bangalore 560012, INDIA

**Author notes:** Department of Biochemistry and Molecular Genetics, Northwestern University Feinberg School of Medicine, Chicago, IL 60611. To whom correspondence should be addressed: Pundi Rangarajan, Department of Biochemistry, Indian Institute of Science, Bangalore 560012, INDIA.

**Keywords:** aldehyde dehydrogenase, Mxr1p, transcriptional regulation, alcohol oxidase, yeast metabolism

## Abstract

The genome of the methylotrophic yeast, *Komagataella phaffii* harbours multiple genes encoding putative alcohol dehydrogenases and aldehyde dehydrogenases (ALDs). Here, we demonstrate that one of the ALDs denoted as ALD-A is essential for ethanol metabolism. A zinc finger transcription factor known as Mxr1p regulates *ALD-A* transcription by binding to Mxr1p response elements (MXREs) in the *ALD-A* promoter. Mutations which abrogate Mxr1p binding to *ALD-A* MXREs *in vitro* abolish transcriptional activation from *ALD-A* promoter *in vivo*. Mxr1p regulates *ALD-A* expression during ethanol as well as methanol metabolism. ALD-A is essential for the utilization of methanol and *Δald-a* is deficient in alcohol oxidase (AOX), a key enzyme of methanol metabolism. AOX protein but not mRNA levels are down regulated in *Δald-a.* ALD-A and AOX localize to cytosol and peroxisomes respectively during methanol metabolism suggesting that they are unlikely interact with each other *in vivo*. This study has led to the identification of Mxr1p as a key regulator of *ALD-A* transcription during ethanol and methanol metabolism of *K. phaffii*. Post-transcriptional regulation of AOX protein levels by ALD-A during methanol metabolism is another unique feature of this study.

## INTRODUCTION

In *S. cerevisiae*, ethanol is metabolized into acetaldehyde, acetate and acetyl-CoA by the activity of alcohol dehydrogenase (ADH), aldehyde dehydrogenase (ALD) and acetyl-CoA synthetase (ACS) respectively. These enzymes are encoded by multiple genes which are well characterized. For example, ADH1 functions as the major fermentative enzyme during glucose metabolism, catalyzing the conversion of acetaldehyde to ethanol (1, 2). ADH2 is the major enzyme in the utilization of ethanol as carbon source its expression is repressed by glucose (3–6). Both ADH1 and ADH2 are cytosolic enzymes. A third isozyme, ADH3, localized in the mitochondria, is a component of the ethanol/acetaldehyde shuttle (7). ADH4 is most closely related to a bacterial, iron-activated alcohol dehydrogenase but requires zinc for its activity like the other *S. cerevisiae* ADH proteins (8–10). *ADH5* codes for a minor activity and has not been well characterized (11).

Aldehyde dehydrogenases (ALDs) play a key role during growth of yeasts on non-fermentable carbon sources by converting acetaldehyde generated from ethanol to acetate. They are also involved in the breakdown of toxic aldehydes accumulated under stress conditions (12). In *S. cerevisiae*, ALDs are encoded by five genes: *ALD1/ALD6 (YPL061w), ALD2 (YMR170c)* and *ALD3 (YMR169c),* which encode cytosolic isoforms, as well as *ALD4 (YOR374w)* and *ALD5 (YER073w),* which encode mitochondrial isoforms (13). Several isoenzymes of *S. cerevisiae* ALD are categorized based on their subcellular locations. *ALD2, ALD3* and *ALD6* encode cytosolic aldehyde hydrogenases whereas *ALD4* and *ALD5* encode mitochondrial enzymes (14). Of these, ALD4 and ALD6 are the two major enzymes. Expression of *ALD6* and *ALD4* is induced and repressed by glucose respectively. Expression of *ALD4* can be induced in the presence of ethanol and acetaldehyde (14). The carbon source-responsive zinc finger transcription factors, Adr1p and Cat8p regulate *S. cerevisiae ADH2* expression by binding to upstream activation sequences 1 and 2 respectively in the *ADH2* promoter (6,15, 16). Adr1p is also required for the activation of genes involved glucose fermentation (e.g., *ALD4* and *ALD6*), glycerol metabolism (e.g., *GUT1* and *GUT2*), fatty acid utilization (e.g., *POX1, FOX2*), and peroxisome biogenesis (e.g., *PEX1*) (17,18).

*Komagataella phaffii* previously known as *Pichia pastoris* is a methylotrophic yeast and the enzymes of methanol metabolism are well characterized (19). Methanol is converted to formaldehyde by the enzyme alcohol oxidase which is encoded by two genes, *AOXI* and *AOX2.* AOX1 is a methanol inducible enzyme and the *AOXI* promoter is widely used for production of recombinant proteins (19). Unlike *S. cerevisiae*, *K. phaffii* is a respiratory yeast capable of growing to high cell densities when cultured in minimal media containing glucose or glycerol. *K. phaffii* can also utilize oleic acid, acetate, sorbitol, ethanol or amino acids as the sole source of carbon. Enzymes of ethanol metabolism as well as the regulatory circuits governing their synthesis are not well characterized in *K. phaffii*. Atleast three *K. phaffii* alcohol dehydrogenases referred to as ADH, ADH2, ADH3 have been reported (20–25). ADH3 is responsible for 92% of the total alcohol dehydrogenase enzyme activity when cells are grown on glucose media and it shares 73.6% and 71.4% amino acid identity with *S. cerevisiae* ADH2 and ADH3, respectively (25). *K. phaffii* ADH3, annotated as mitochondrial alcohol dehydrogenase is likely to be a cytosolic ADH as it does not possess a mitochondrial targeting sequence (22, 25). *K. phaffii ADH3* promoter has been exploited for production of recombinant proteins (26). Deletion of *K. phaffii ADH2* does not affect the growth of *K. phaffii* on glucose or ethanol containing media (21). Acetyl-CoA, a downstream metabolite of ethanol was shown to repress enzymes of methanol metabolism when cultured in presence of ethanol (27).

*K. phaffii* is cultured in minimal medium containing yeast nitrogen base (YNB) or nutrient-rich medium containing yeast extract, peptone (YP) supplemented with different sources of carbon such as glucose, glycerol, methanol, ethanol, acetate, glutamate etc. Several transcription factors such as Mxr1p, Rop1p, Trm1p, Nrg1p, Mit1p, Mig1p and Mig2p (19, 28–38) regulate the expression of target genes of specific metabolic pathways of *K. phaffii.* In addition to conventional sources of carbon such as glucose and glycerol, *K. phaffii* can utilize amino acids such as glutamate as the sole source of carbon and hence can be cultured in YNB medium containing 2% glutamate (YNB+Glu) or YP medium in which amino acids derived from peptone serve as the source of carbon (38). Mxr1p regulates the expression of several enzymes required for glutamate utilization in cells cultured in YP or YNB+Glu (38). Mxr1p regulates the expression of *ACS1* encoding acetyl-CoA synthetase 1 during acetate metabolism (37). The target genes of Mxr1p harbour Mxr1p response elements (MXREs) in their promoters to which Mxr1p binds and activates transcription (28–31,37,38). Transcription factors regulating ethanol metabolism of *K. phaffii* have not been identified. In *S. cerevisiae and Kluyveromyces lactis,* expression of *ADH2* is regulated by Adr1p considered as the homologue of Mxr1p (15, 39, 40). This study was initiated to examine the role of Mxr1p in the regulation of ethanol metabolism and identify its target genes. Here, we demonstrate that Mxr1p is essential for ethanol metabolism and unlike *S. cerevisiae* Adr1p, Mxr1p does not regulate the expression of *ADH* genes of *K. phaffii.* Instead, it regulates the transcription of *ALD-A* encoding one of the four putative ALDs of *K. phaffii* by binding to MXREs in the *ALD-A* promoter. Interestingly, Mxr1p regulates *ALD-A* expression during methanol metabolism as well and the ALD-A has a novel role as regulator of AOX, a key enzyme of methanol metabolism.

## RESULTS

### Putative alcohol dehydrogenases (ADHs) and aldehyde dehydrogenases (ALDs) of *K. phaffii*

*K. phaffii* genome contains multiple genes with varying degrees of homology to *S. cerevisiae ADH* and *ALD* genes. While these are annotated as *K. phaffii ADH* and *ALD*s, whether they are functional homologues of *S. cerevisiae ADH* and *ALD*s is not clear. Since the exact function of several *K. phaffii* ADHs and ALDs is not known, they were designated using the alphabets A,B,C,D instead of Arabic numerals (1,2,3,4 etc.,) to distinguish them from *S. cerevisiae* enzymes as indicated in Table 1. *K. phaffii* strains used in this study are listed in Table 2. *K. phaffii* strains are cultured in a medium containing 0.17% yeast nitrogen base (YNB) and 2% glucose (YNBD), 1% ethanol (YNBE) or 1% methanol (YNBM).

**Table 1.** *K. phaffii* alcohol dehydrogenase (ADHs) and aldehyde dehydrogenases (ALDs).

| *K. phaffii* gene | Designation in this study | References |
| --- | --- | --- |
| **ADH:** |  |  |
| *PAS_chr3_0006* | ADH-A (XP_002492217.1; XM_002492172.1) | 44 |
| *PAS_chr1-1_0357* | ADH-B (XP_002490014.1; XM_002489969.1) | 44 |
| *PAS_chr2-1_0472* | ADH-C* (XP_002491382.1; XM_002491337.1) | 26, 27, 45, 46 |
| *PAS_chr4_0576* | ADH-D@ (XP_002494014.1; XM_002493969.1) | 45 |
| *PAS_chr3_0349* | ADH-E (XP_002492569.1; XM_002492524) | 44 |
| *PAS_chr2-1_0313* | ADH-F (XP_002491208.1; XM_002491163.1) | 44 |
| **ALD:** |  |  |
| *PAS_chr4_0043* | ALD-A (XP_002493450.1) (XM_002493405.1) | This study |
| *PAS_chr3_0987* | ALD-B# (XP_002493229.1) (XM_002493184.1) | 45 |
| *PAS_chr2-1_0853* | ALD-C+ (XP_002491418.1) (XM_002491373.1) | 27, 45 |
| *PAS_chr2-1_0453* | ALD-D\$ (XP_002491360.1) (XM_002491315.1) | 45 |

GenBank accession numbers are shown in parentheses.
\*denoted as ADH2 (27, 46, 45) or ADH3 (26).
@denoted as ADH6 (45)

**Table 2.** *K. phaffii* strains used in this study.

| Strain | Genotype | References |
| --- | --- | --- |
| *GS115* | *his4* | 28 |
| *KM71 (Δaox1)* | *Δaox1::ARG4 his4 arg4* | ThermoFisher |
| *Δmxr1* | *GS115, Ppmxr1Δ::His4* | 28 |
| *Δmxr1* | *GS115, Ppmxr1Δ::Zeocin<sup>r</sup>* | 37 |
| *Δald-a* | *GS115, Ppald-a Δ::Zeocin<sup>r</sup>* | This study |
| *Δald-d* | *GS115, Ppald-d Δ::Zeocin<sup>r</sup>* | This study |
| *GS115-ALD-A<sup>Myc</sup>* | *GS115, His4<sup>+</sup>::(P<sub>ALD-A</sub>PpALD-A<sup>Myc</sup>)* | This study |
| *Δmxr1-ALD-A<sup>Myc</sup>* | *Δmxr1, His4<sup>+</sup>::(P<sub>ALD-A</sub>PpALD-A<sup>Myc</sup>)* | This study |
| *GS115-P<sub>ALD-A</sub>-GFP* | *GS115, His4<sup>+</sup>::(P<sub>ALD-A</sub>GFP)* | This study |
| *Δmxr1-P<sub>ALD-A</sub>-GFP* | *Δmxr1, His4<sup>+</sup>::(P<sub>ALD-A</sub>GFP)* | This study |
| *GS115-ALD-A-M1<sup>GFP</sup>* | *GS115, His4<sup>+</sup>::(P<sub>ALD-A-M1</sub>GFP)* | This study |
| *GS115-ALD-A-M2<sup>GFP</sup>* | *GS115, His4<sup>+</sup>::(P<sub>ALD-A-M2</sub>GFP)* | This study |
| *GS115-ALD-A-M2,3<sup>GFP</sup>* | *GS115, His4<sup>+</sup>::(P<sub>ALD-A-M2</sub>GFP)* | This study |
| *GS115-ALD-A-M3<sup>GFP</sup>* | *GS115, His4<sup>+</sup>::(P<sub>ALD-A-M3</sub>GFP)* | This study |
| *GS115-ALD-A-M<sup>\*GFP</sup>* | *GS115, His4<sup>+</sup>::(P<sub>ALD-A-M\*</sub>GFP)* | This study |
| *GS115-P<sub>AOX1</sub>-GFP* | *GS115, His4<sup>+</sup>::(P<sub>AOX1</sub>GFP)* | This study |
| *Δald-a-P<sub>AOX1</sub>-GFP* | *Δald-a, His4<sup>+</sup>::(P<sub>AOX1</sub>GFP)* | This study |

### Transcriptional regulation of *ALD-A* during ethanol and methanol metabolism by Mxr1p

The zinc finger transcription factor Mxr1p has emerged as a key regulator of multiple metabolic pathways of *K. phaffii*. However, its role in the regulation of genes of ethanol metabolism is not known. We therefore examined whether Mxr1p is essential for ethanol metabolism by analyzing the growth of *K. phaffii GS115* and *Δmxr1* strains in YNBE. The results indicate that *Δmxr1* exhibits retarded growth in YNBE (Fig. 1A). Utilization of ethanol as a source of carbon involves its sequential conversion into acetaldehyde, acetate and acetyl-CoA by ADH, ALD and ACS respectively (Fig. 1B). *K. phaffii* genome contains multiple *ADH*s and *ALD*s (Table 1). To examine whether genes are expressed, we examined their mRNA levels by qPCR in cells cultured in YNBD and YNBE. The results indicate that *ADH*s and *ALD*s are actively transcribed during glucose and/or ethanol metabolism (Fig. 1C).

**Fig. 1.**
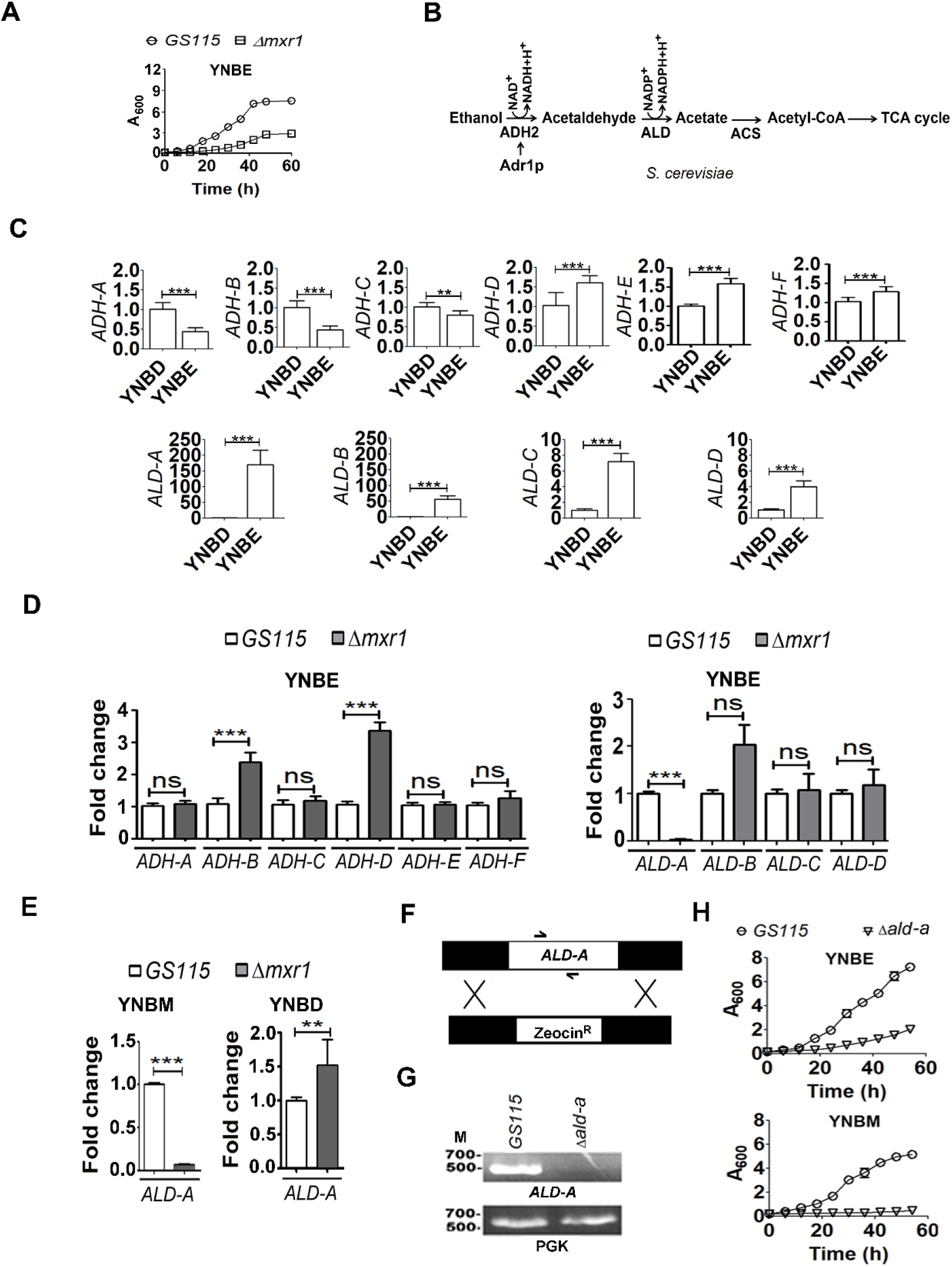
Identification of *ALD-A* as the target of Mxr1p and analysis of its function during ethanol and methanol metabolism. **A.** Analysis of growth of *GS115* and *Δmxr1* cultured in YNBE and YPE. **B.** Schematic representation of ethanol utilization pathway of *S. cerevisiae*. *S. cerevisiae* genome encodes NAD^+^ as well as NADP^+^ ALDs (47). **C.** Quantitation of mRNAs encoding ADHs and ALDs of *K. phaffii* by qPCR in *GS115* cultured in YNBD and YNBE. **D.** Quantitation of mRNAs encoding ADHs and ALDs of *K. phaffii* by qPCR in *GS115* and *Δmxr1* cultured in YNBE. Error bars in each figure indicate S.D. Data is from three biological replicates (n=3). In the graphs, P value summary is mentioned on the bar of each figure. *p<0.05, **p<0.005, ***p<0.0005, ns, not significant. Student’s paired or unpaired t-test was done. **E.** Quantitation of *ALD-A* mRNA levels by qPCR in *GS115* and *Δmxr1* cultured in YNBM and YNBD. Error bars in each figure indicate S.D. Data is from three biological replicates (n=3). In the graphs, P value summary is mentioned on the bar of each figure. *p<0.05, **p<0.005, ***p<0.0005, ns, not significant. Student’s paired or unpaired t-test was done. **F.** Strategy for the generation of *Δald-a.* **G.** Confirmation of deletion of *ALD-A* by PCR. *ALD-A* was amplified from genomic DNA by PCR using gene-specific primers. **H.** Analysis of growth of *GS115* and *Δmxr1* cultured in YNBE and YNBM.

In *S. cerevisiae,* Adr1p, often considered as the homologue of *K. phaffii* Mxr1p (15,28), regulates the expression of *ADH2,* which encodes aldehyde dehydrogenase 2, the first enzyme of ethanol metabolism (Fig. 1B) (28). We examined whether Mxr1p is required for the expression of any of the *ADH*s in *K. phaffii* by measuring their mRNA levels by qPCR in *GS115* and *Δmxr1* cultured in YNBE. The results indicate that none of the *ADH* mRNAs are down regulated in *Δmxr1*(Fig. 1D). Interestingly, *ADH-B* and *ADH-D* mRNAs are upregulated in *Δmxr1* (Fig. 1D) although the significance of their upregulation is not clear at his juncture. Analysis of mRNA levels of *ALD*s indicate that *ALD-A* is downregulated in *Δmxr1* cultured in YNBE (Fig. 1D). Interestingly, *ALD-A* mRNA levels are also down regulated in *Δmxr1* cultured in YNBM but not YNBD (Fig. 1E). To understand ALD-A function during ethanol and methanol metabolism, *K. phaffii Δald-a* carrying a deletion of *ALD-A* was generated (Fig. 1F,G) and its growth was examined in cells cultured in YNBE and YNBM. *Δald-a* exhibited impaired growth when cultured in YNBE as well as YNBM (Fig. 1H) indicating that ALD-A is essential for the metabolism of not only ethanol but also methanol. To further confirm regulation of ALD-A expression by Mxr1p, ALD-A was expressed as a Myc-tagged protein (ALD-A^Myc^) from *ALD-A* promoter and ALD-A protein levels were examined by western blot analysis using anti-Myc antibodies. ALD-A^Myc^ levels in *Δmxr1* were lower than those in *GS115* cultured in YNBE and YNBM but not YNBD (Fig. 2A). GFP was expressed from *ALD-A* promoter and its expression was examined by western blotting using anti-GFP antibodies. The results indicate that GFP levels are significantly lower in *Δmxr1* than *GS115* cultured in YNBE and YNBM (Fig. 2B). These results confirm that Mxr1p is a key regulator of *ALD-A* expression. Immunofluorescence studies indicate that ALD-A^Myc^ predominantly localizes to the cytoplasm of cells cultured in YNBE and YNBM (Fig. 2C).

**Fig. 2.**
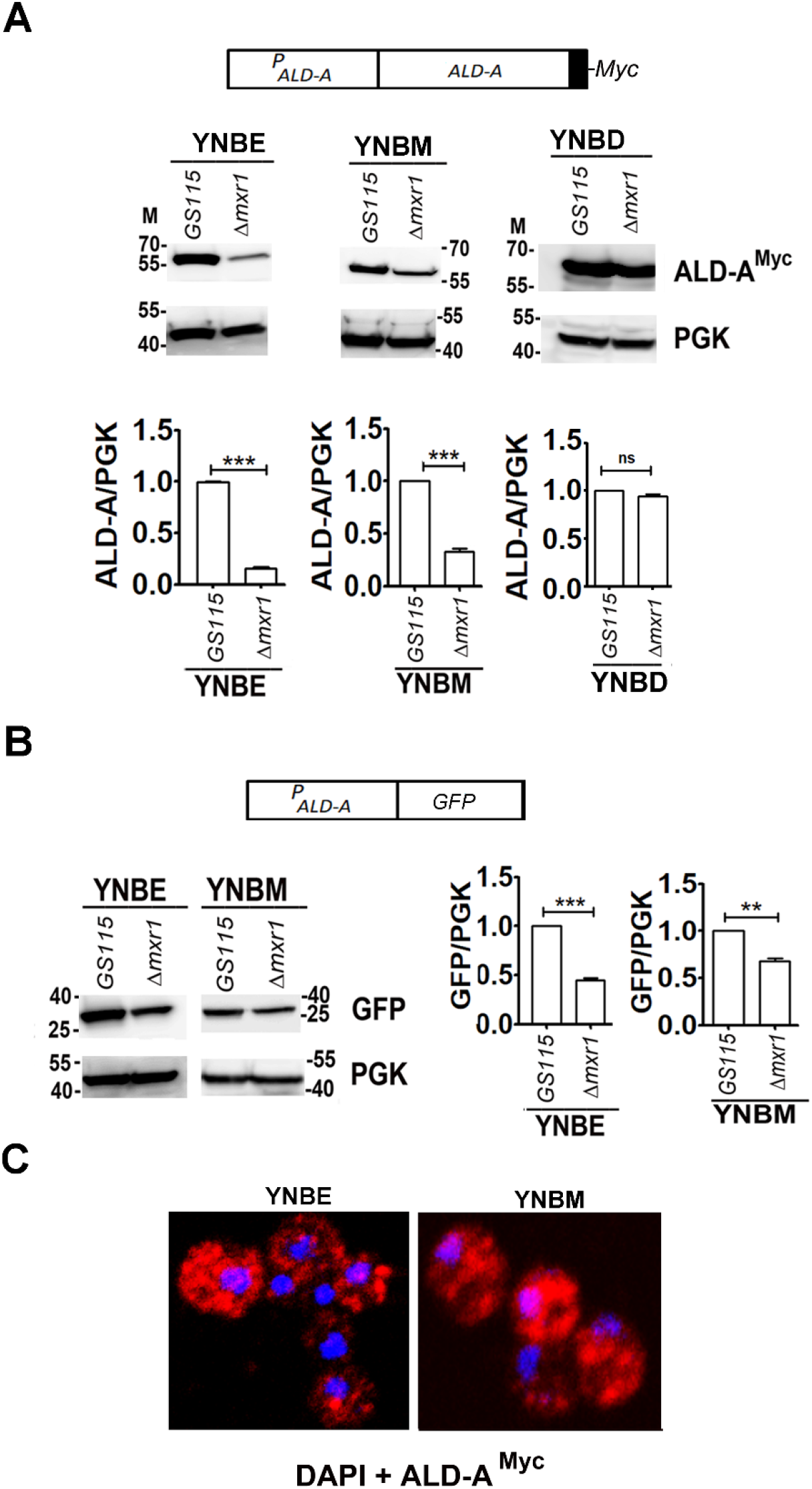
Confirmation of Mxr1p as a regulator of ALD-A expression. **A.** Analysis of ALD-A^Myc^ levels in *GS115* and *Δmxr1* in cells cultured in YNBE, YNBM and YNBD by western blotting using anti-Myc antibodies. M, protein molecular weight markers (kDa). Quantitation of the data is presented. The intensity of individual bands was quantified and expressed as arbitrary units± S.D. relative to controls. Data is from three biological replicates (n=3). **B.** Schematic representation of *P*_*ALD-A*_*GFP* and analysis of GFP levels in *GS115* and *Δmxr1* cultured in YNBE and YNBM by western blotting using anti-GFP antibodies. PGK was used as loading control. M, protein molecular weight markers (kDa). Quantitation of the data presented in B is also shown. The intensity of individual bands was quantified and expressed as arbitrary units± S.D. relative to controls. Data is from three biological replicates (n=3). **C.** Subcellular localization of ALD-A^Myc^ as analyzed by immunofluorescence using confocal microscopy. Mouse anti-Myc antibodies and Alexa Flour 488-conjugated, donkey anti-mouse antibodies were used. DAPI was used to stain the nucleus.

### Identification of Mxr1p response elements (MXREs) in *ALD-A* promoter (*P*_*ALD-A*_)

Mxr1p regulates the expression of target genes by binding to MXREs in their promoters which bear the consensus sequence 5’ CYCCNY 3’ (29,37,38). Analysis of the nucleotide sequence of 1.0 kb *ALD-A* promoter indicated the presence of three putative MXREs designated as MXRE1, MXRE2 and MXRE3 (Fig. 3A). Since point mutations within the 5’ CYCCNY 3’ motif of *AOXI* MXREs abrogate Mxr1p binding (29), similar mutations were introduced within the 5’ CYCCNY 3’ motif of the putative *ALD-A* MXREs and these were designated as MXRE1-M, MXRE2-M and MXRE3-M (Fig. 3B). The effect of these mutations on the binding of recombinant Mxr1p containing 150 N-terminal amino acids including the zinc finger DNA binding domain (Mxr1p^N150^) (38) to *ALD-A* MXREs was examined in an electrophoretic mobility shift assay. The results indicate that Mxr1p^N150^ binds to wild type but not the mutant MXREs (Fig. 3C). We examined the ability of Mxr1p to activate transcription from *ALD-A* promoter containing wild type and mutant MXREs. The gene encoding GFP was cloned downstream of 1.0 kb *ALD-A* promoter containing wild type and mutant MXREs (Fig. 4A), transformed into *GS115* and GFP expression was examined in cells cultured in YNBE and YNBM. The results indicate that point mutation in MXRE1, 2 or 3 resulted in only a marginal reduction or no reduction in GFP expression (Fig. 4B). However, GFP expression was drastically reduced when all the three or two MXREs were mutated (Fig. 4B-D). These results are summarized in Fig. 4E.

**Fig. 3.**
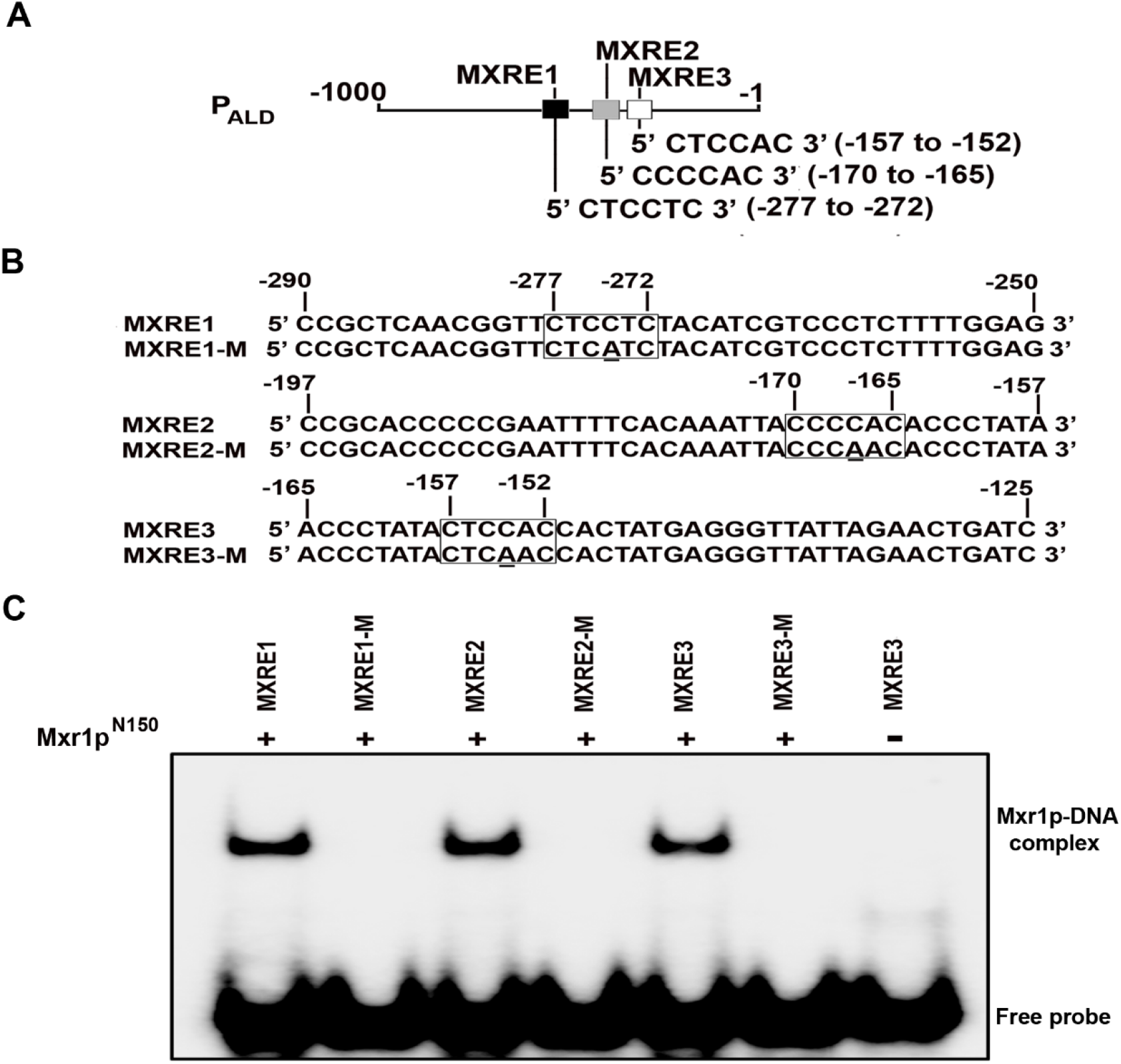
Identification of MXREs in *ALD-A* promoter. **A.** Schematic representation of position of putative MXREs in *ALD-A* promoter. **B.** Nucleotide sequence of oligonucleotides used in EMSA. MXREs are boxed and point mutations within MXREs are underlined. **C.** Analysis of recombinant Mxr1p^N150^ binding to radiolabeled *ALD-A* promoter regions by EMSA. Point mutations within MXREs abrogate Mxr1p binding.

**Fig. 4.**
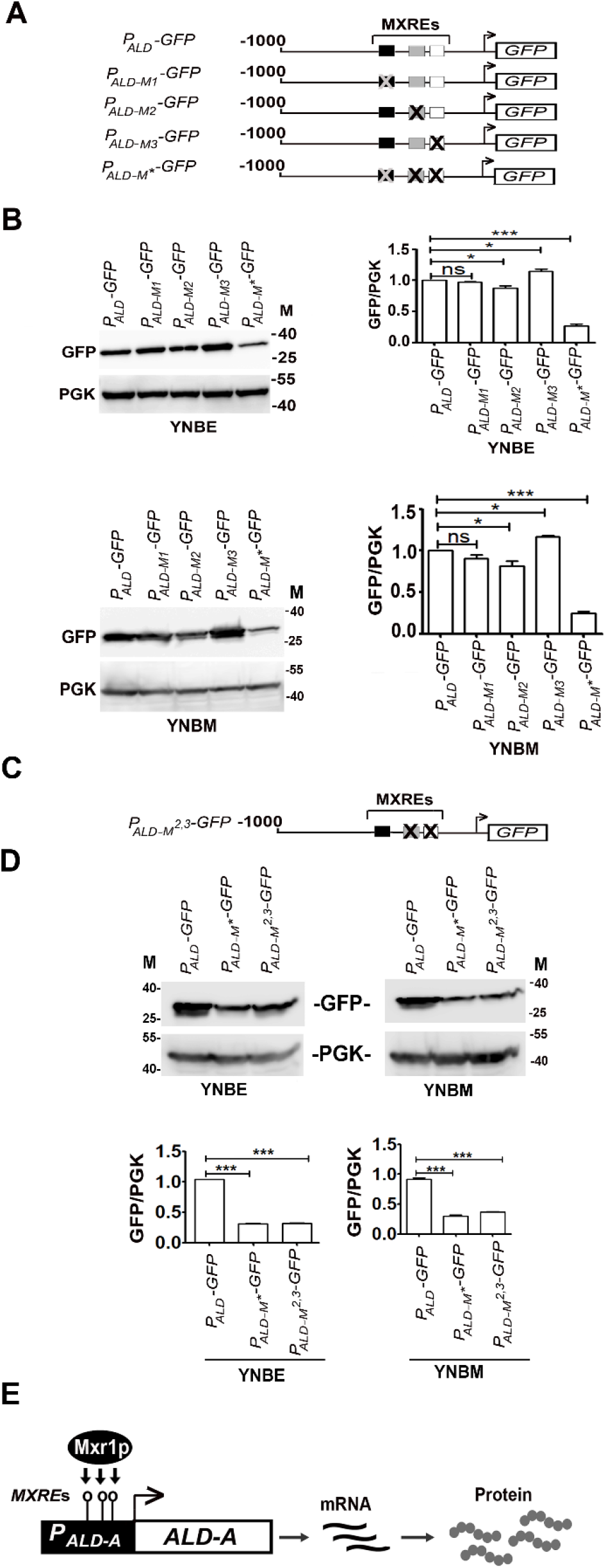
Analysis of function of MXREs of ALD-A promoter *in vivo*. **A.** Schematic representation of different *P*_*ALD-A*_*GFP* constructs. **B.** Analysis of GFP expression from *ALD-A* promoter containing wild type or mutant MXREs by western blotting using anti-GFP antibodies in cells cultured in YNBE and YNBM. PGK was used as loading control. M, protein molecular weight markers (kDa). Quantitation of the western blot data is also shown. The intensity of individual bands was quantified and expressed as arbitrary units± S.D. relative to controls Data is from three biological replicates (n=3). **C.** Schematic representation of *P*_*ALD-A*_*GFP* construct carrying two mutant MXREs. **D.** Analysis of GFP expression from *ALD-A* promoter containing wild type or mutant MXREs by western blotting using anti-GFP antibodies in cells cultured in YNBE and YNBM. PGK was used as loading control. M, protein molecular weight markers (kDa). Quantitation of the western blot data is also shown. The intensity of individual bands was quantified and expressed as arbitrary units± S.D. relative to controls. Data is from three biological replicates (n=3). **E.** Schematic representation of regulation of *ALD-A* by Mxr1p. during ethanol and methanol metabolism.

### An essential role for ALD-A during methanol metabolism

The inability of *Δald-a* to grow in YNBM suggested that ALD-A is essential for methanol metabolism as well. To gain further insights, we first examined the protein profile of cell lysates of *GS115* and *Δald-a* cultured in YNBM. AOX is the most abundant protein in cells metabolizing methanol and the protein band can be readily visualized in SDS polyacrylamide gels stained with Coomassie Brilliant Blue R. A protein band corresponding to the molecular weight of AOX was present at high levels in *GS115* but not *Δald-a* cultured in YNBM (Fig. 5A). To confirm AOX down regulation in *Δald-a*, western blot analysis was carried out with anti-AOX antibodies. The results revealed that AOX protein levels are lower in *Δald-a* than *GS115* (Fig. 5B). A decrease in AOX protein levels in *Δald-a* was not accompanied by a decrease in *AOX1* mRNA levels (Fig. 5C) suggesting that ALD-A-mediated decrease in AOX protein levels may involve a post-transcriptional mechanism. *GS115-P*_*AOX1*_-*GFP* strain contains *P*_*AOXI*_*-GFP* expression cassette integrated at the *His4* locus (Table 2) and therefore expresses AOX1 protein from the native *AOX1* promoter as well as GFP from the 1.0 kb *AOXI* promoter at *His4* locus. Lysates were prepared from this strain cultured in YNBM, AOX and GFP protein levels were quantitated by western blot analysis using anti-AOX and anti-GFP antibodies respectively. Only AOX but not GFP protein levels were reduced in *Δald-a* (Fig. 5D) indicating that only AOX protein but not GFP expressed from *AOX1* promoter is subjected to ALD-A mediated down regulation. AOX is known to localize to peroxisomes during methanol metabolism (41) and this was confirmed by immunofluorescence using anti-AOX antibodies in *GS115* cultured in YNBM (Fig. 5E). Under similar culture conditions, ALD-A^Myc^ localizes to cytosol (Fig. 5E). ALD-A localizes to cytosol in cells cultured in YNBE as well (Fig. 2C). Thus, AOX and ALD-A localize to two distinct sub-cellular compartments and therefore unlikely to interact with each other *in vivo*.

**Fig. 5.**
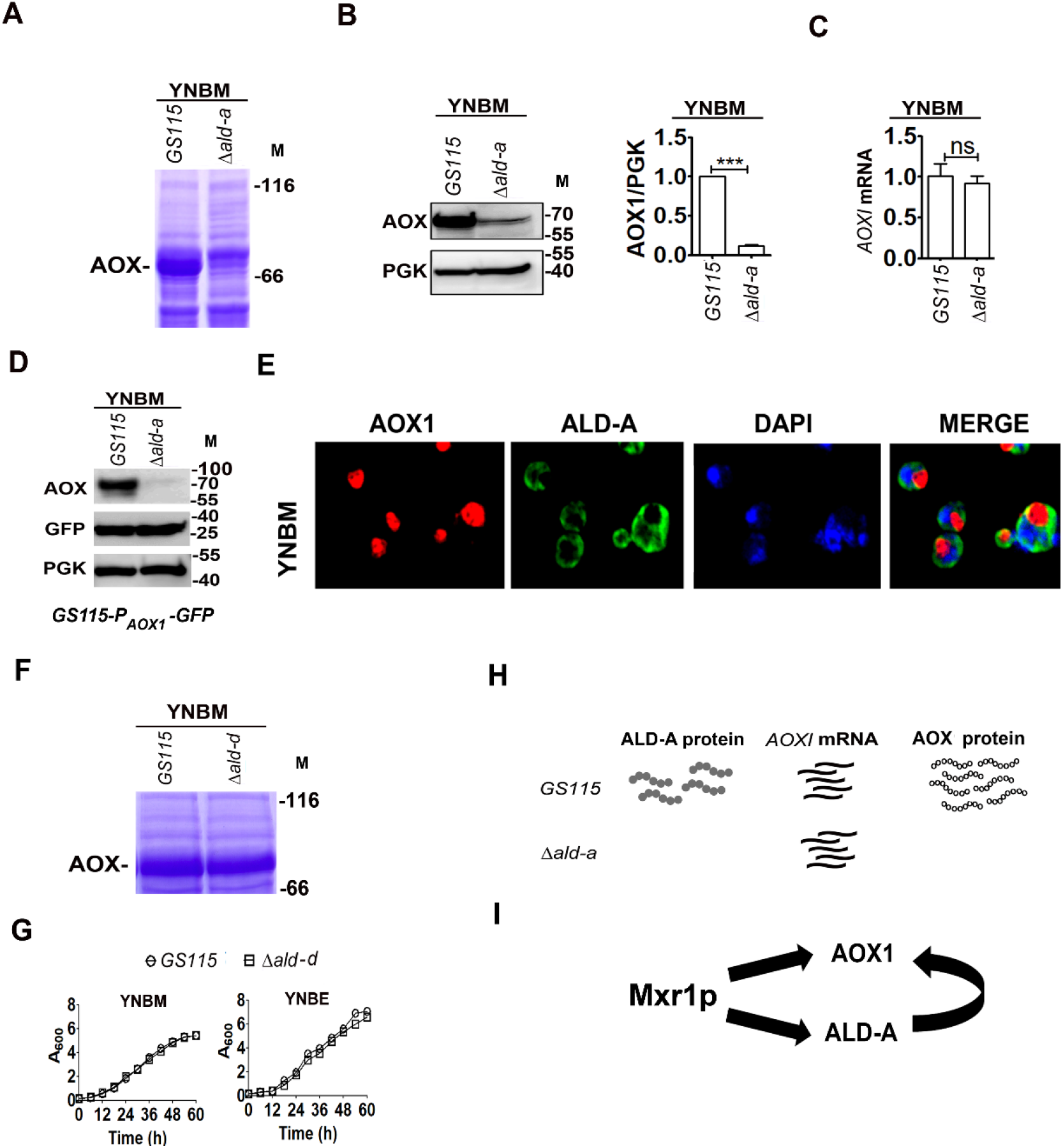
Regulation of AOX protein levels by ALD-A in cells cultured in YNBM. **A.** Analysis of AOX protein levels in *GS115* and *Δald-a* cultured in YNBM by SDS-PAGE. Gels were stained with Coomassie Brilliant blue R. Protein molecular weight markers (kDa) are indicated (M). **B.** Analysis of AOX protein levels in *GS115* and *Δald-a* cultured in YNBM by western blotting. Anti-AOX antibodies were used. M, protein molecular weight markers. Quantification of intensity of bands is shown. Data is from three biological replicates (n=3). **C.** Quantification of *AOX1* mRNA levels by qPCR. Error bars indicate S.D. n=3. ns, not significant. **D.** Quantification of AOX and GFP expressed from *AOXI* promoter by western blotting using anti-AOX and anti-GFP antibodies respectively. PGK was used as loading control. Lysates were prepared from *GS115P*_*AOX*_-*GFP* strain in which AOX protein is synthesized from the *AOX1* and *AOX2* loci while GFP is synthesized from 1.0 kb *AOX* promoter of the *P*_*AOX1*_-*GFP* expression cassette integrated at *His4* locus. **E.** Subcellular localization of ALD-A^Myc^ and AOX as analyzed by immunofluorescence by confocal microscopy using rabbit anti-Myc antibodies and mouse anti-AOX antibodies respectively. DAPI was used to stain the nucleus. **F.** Analysis of AOX protein levels in *GS115* and *Δald-d* cultured in cells cultured in YNBM and YPM by SDS-PAGE. Gels were stained with Coomassie Brilliant blue R. M, protein molecular weight markers (kDa). **G.** Analysis of growth of *GS115* and *Δald-d* cultured in YNBE and YNBM. **H.** Schematic representation of regulation of AOX protein levels by ALD-A during methanol metabolism. **I.** Schematic representation of regulation of AOX expression during methanol metabolism. Mxr1p regulates *AOX1* and *ALD-A* expression at the transcriptional level and ALD-A in turn regulates AOX protein levels at the post-transcriptional level.

*P. pastoris* genome encodes four ALDs and we examined the effect of deletion of *ALD-D* on AOX protein levels as well as growth. The results indicate that in AOX protein level in *Δald-d* is comparable to that of *GS115* (Fig. 5F). Further, deletion of *ALD-D* had no effect on the growth of cells cultured in YNBM and YNBE (Fig. 5G). These results are summarized in Fig. 5H, I.

## DISCUSSION

In this study, we demonstrate that *K. phaffii* Mxr1p binds to the MXREs present in the promoter of *ALD-A* encoding an aldehyde dehydrogenase and activates its transcription during the metabolism of ethanol and methanol. This is the first report on the trans-activation of a gene essential for ethanol metabolism by Mxr1p. Thus far, studies have only focused on the repression of genes of methanol utilization pathway such as *AOX1* during ethanol metabolism. It was suggested that acetyl-CoA synthesized during ethanol metabolism may be utilized for the acetylation of histones or other TFs which may repress the expression of methanol-inducible genes (Karaoglan *et al*., 2016b). In another study, phosphorylation of serine 215 residue of Mxr1p was shown to be involved in the transcriptional repression of *AOXI* during ethanol metabolism (Parua *et al*., 2012). However, the ability of Mxr1p to activate the expression of gene(s) essential for ethanol metabolism has not been investigated. Here, we demonstrate that Mxr1p activates *ALD-A* transcription during the metabolism of not only ethanol but also methanol. *ALD-A* is a key target of Mxr1p in cells cultured in YNBE and YNBM. *ALD-A* promoter (−1.0 kb) harbours three MXREs to which Mxr1p binds *in vitro* and activates transcription *in vivo*. Point mutations known to abrogate Mxr1p binding to *AOX1 MXRE*s (Kranthi *et al*., 2009) abolish Mxr1p binding to *ALD-A MXRE*s as well. Analysis of GFP expression from *ALD-A* promoters containing wild type or mutant MXREs indicate that atleast two MXREs are required for Mxr1p-mediated trans-activation of *ALD-A* promoter *in vivo*. It is pertinent to note that the promoter of *ACS1* encoding acetyl-CoA synthetase 1 consists of two MXREs, both of which are required for Mxr1p-mediated trans-activation during acetate metabolism (37).

Of the four *ALD* genes of *P. pastoris*, we have examined the function of *ALD-A* and *ALD-D* in this study. While deletion of ALD-A results in impaired growth in media containing ethanol and methanol, deletion of *ALD-D* had no significant effect on the growth of cells metabolizing ethanol and methanol. Studies aimed at understanding the role of ALD-A during methanol metabolism indicate that ALD-A is essential for the utilization of methanol and it is required for the maintenance of normal levels of AOX protein in cells cultured in YNBM and YPM. ALD-A is neither a transcription factor nor it localizes to nucleus and therefore unlikely to regulate *AOX* gene transcription. This is substantiated by the fact that only AOX protein but not mRNA levels are down regulated in *Δald-a*. Further, only AOX but not GFP expressed from *AOX1* promoter is down regulated in *Δald-a* indicating that *AOX1* promoter has no role in the ALD-A mediated down regulation of AOX. This is not a general property of all aldehyde dehydrogenases since AOX protein reduction is not seen in *Δald-d.* ALD-A and AOX localize to two different cellular compartments and therefore the down regulation of AOX protein is not due to direct protein-protein interactions. Formaldehyde generated from AOX-catalyzed reaction is converted to formate by formaldehyde dehydrogenase encoded by *FLD1* and therefore a catalytic role for ALD-A in methanol metabolism is unlikely. Thus, the exact mechanism by which AOX protein levels are depleted in *Δald-a* needs further investigations.

This study has led to the identification of Mxr1p as a key regulator of *ALD-A* which is essential for the metabolism of ethanol as well as methanol. Thus far, MXREs have been identified in genes involved in the metabolism of methanol, ethanol, glycerol, acetate and amino acids (28–34, 37, 38, 42, 45, this study) and Mxr1p has achieved a unique status as a global regulator of multiple metabolic pathways in *K. phaffii* (Fig. 6).

**Fig. 6.**
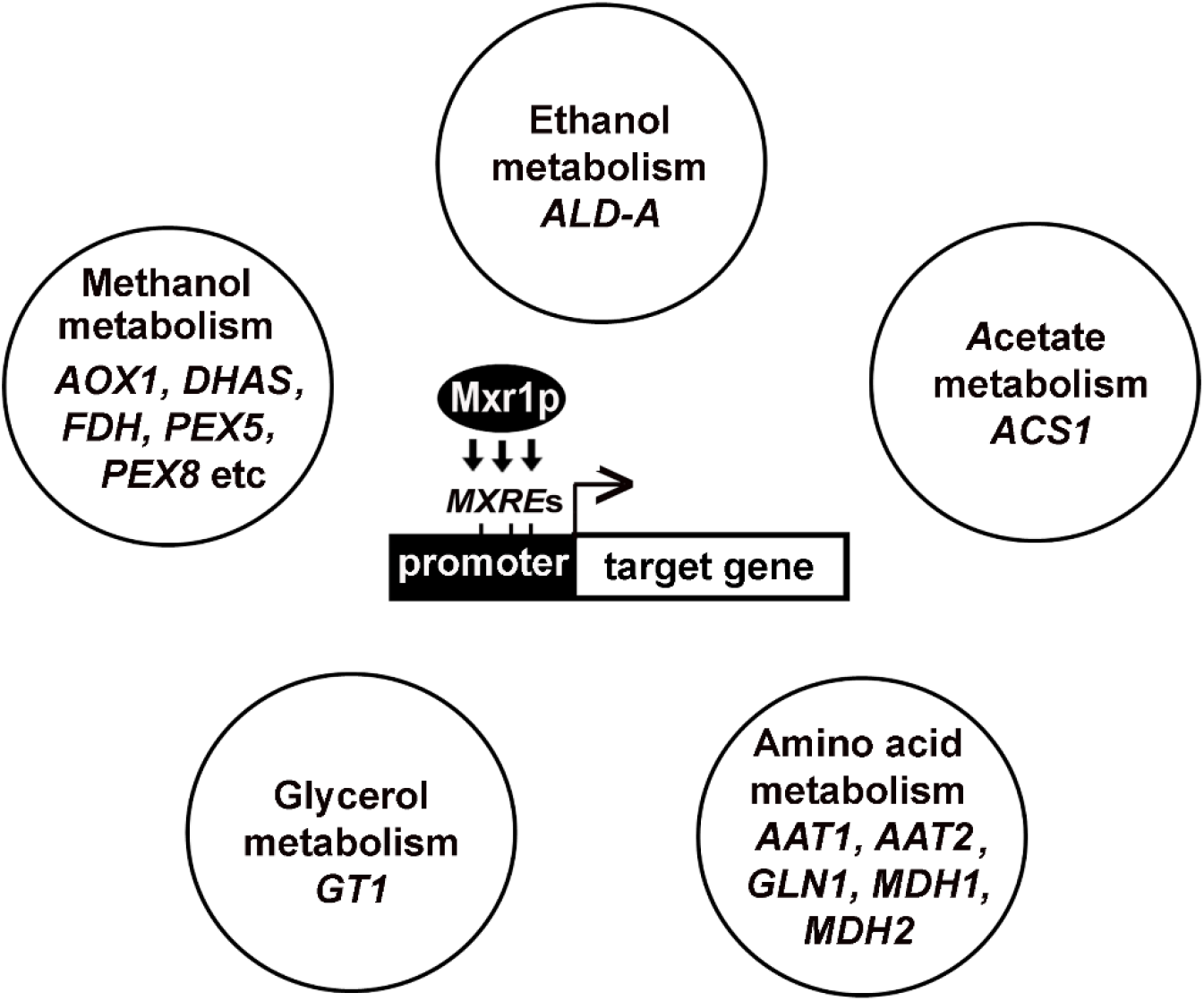
Schematic representation depicting Mxr1p as a global regulator of multiple metabolic pathways of *K. phaffii*. Target genes of Mxr1p containing MXREs in their promoters are shown (28–34, 37, 38, 42, 45, this study).

## EXPERIMENTAL PROCEDURES

### Growth media and culture conditions

Wild type *(GS115, His*^−^) strain of *K. phaffii* was a kind gift of James Cregg (28). *K. phaffii KM71* strain (referred to as *ΔAOX1* in this study) was purchased from Thermo Fisher Scientific. Cells were cultured in YP (1 % yeast extract, 2 % peptone) medium consisting of 2 % dextrose (YPD) or minimal media containing 0.17 % yeast nitrogen base without amino acids and 0.5 % ammonium sulfate (YNB) and 2 % dextrose (YNBD), 1 % methanol (YNBM) or 1 % ethanol (YNBE). All components of cell culture media were purchased from Becton and Dickinson (BD) Biosciences. Yeast transformations were performed using Gene Pulser (Bio-Rad) as per manufacturer’s instructions. For the isolation of recombinant plasmids, DH5*α* strain of *Escherichia coli* was used. Expression of recombinant proteins was carried out using *BL21(DE3)* strain of *E. coli*. PEG method was used to prepare chemically component bacterial cells for transformation.

### Antibodies and other reagents

Anti-AOX1 polyclonal antibodies and anti-phosphoglycerate kinase (PGK) polyclonal antibodies have been described (43). Other antibodies used are: anti-MYC (Merck Millipore, OP-10), anti-GFP (Santa Cruz, SC-9996), Donkey, anti-mouse Alexa Flour 555 and 488 (A31570, A11001, Thermo Fisher). Western blotting was performed as described (43) and ImageJ software was applied to quantify all western blot data. Band intensity of the protein of interest was normalized to that of the loading control, PGK. Restriction enzymes, Taq DNA polymerase, and T4 DNA ligase were purchased from New England Biolabs (Frankfurt, Germany). Oligonucleotides were purchased from Sigma-Aldrich, India. Nucleotide sequence of primers used in qPCR and RT-PCR reactions will be provided on request.

### Subcellular localization studies

Localization of AOX and ALD-A^Myc^ by immunofluorescence using anti-AOX and Anti-MYC antibodies was carried out essentially as described (43). Fluorescent microscope (Leica DMLA) or confocal microscope Zeiss LSM 880 was used for visualization and capture of images.

### Quantitative real time PCR (qPCR)

*K. phaffii* cells were cultured for 12-14 h and total RNA was isolated using RNA isolation kit (Cat. # Z3100, Promega) as per manufacturer’s instructions. cDNA was prepared and qPCR was carried out using iQ SYBR Green super mix and iQ5 multicolour real time PCR thermal cycler (Bio-Rad). The levels of mRNA expression relative to GS115 was normalized to 18S rRNA. The comparative Ct method for relative quantification (ΔΔCt method), which describes the change in expression of the target genes in a test sample relative to a calibrator sample, was used to analyze the data.

### Statistical analysis

This was carried out essentially as described (43). Briefly, statistical tests such as student’s t-test, one-way analysis of variance (*ANOVA*) followed by Tukey’s multiple comparison were carried out using GraphPad Prism 5 software. Data are represented as mean ± S.D. *P* value summary is mentioned on the bar of each figure where * *P<0.05*; ** *P<0.005*; *** *P<0.0005*, ns, not significant.

### Mass spectrometry

The protein bands of interest were excised precisely from SDS polyacrylamide gel and subjected to in-gel trypsin digestion using sequencing grade trypsin (Promega, USA). MALDI TOF was performed in HCT Ultra PTM Discovery System (ETD II-Bruker Daltonics) with 1100 series HPLC (Agilent). MASCOT protein mass fingerprint software with NCBI non-redundant database was used to identify target proteins.

### Electrophoretic mobility shift assay

This was carried out with radiolabeled *ALD-A* promoter regions and recombinant Mxr1p consisting of N-terminal 150 amino acids (Mxr1p^N150^) as described (29).

### *K. phaffii* strains

#### Δmxr1

*Δmxr1* strains in which *MXR1* was disrupted by *HIS4* or zeocin resistance cassette (*Zeo*^*R*^) have been described (28, 37).

#### Δald-a, Δald-d

A 0.911 kb *ALD-A* promoter region was amplified by PCR from *K. phaffii* genomic DNA using primer pair 5’-GGACTGTTCAATTTGAAGTCGATGCTGACG-3’ and 5’-GCTATGGTGTGTGGGGGATCCGCACACGATCCCTTGGGAACTTGCGGTGG-3’ (962 to 984 bp of *GAPDH* promoter in reverse complement, −89 to −114 bp of *ALD-A* promoter in reverse complement). In another PCR, 1.2 kb of zeocin expression cassette was amplified by PCR from pGAPZA vector using the primer pair 5’- CCACCGCAAGTTCCCAAGGGATCGTGTGCGGATCCCCCACACACCATAGC-3’ (−89 to −114 bp of *ALD-A* promoter, +962 to +984 bp of *GAPDH* promoter) and 5’GGAGTGTAAG CAATTCTGATAGCCTTGTGCCACATGTTGGTCTCCAGCTTG-3’ (+1467 bp to +1493 bp in reverse complement of 3’-flanking region of *ALD-A*, +2137 to +2159 bp in reverse complement of *GAPDH* promoter). In the third PCR, 872 bp of the 3’-flanking region of *ALD-A* was amplified using the primer pair 5’-CAAGCTGGAGA CCAACATGTGAGCACAGGCTATCAGAATTGCTTACACTCC-3’ (+2137 to +2159 bp of *GAPDH* promoter, +1467 to +1493 bp of region of *ALD-A* gene) and 5’- GGAACTGGAG GCTTCCGCAGCAAACTCTC-3’(+2352 to +2381 bp in the reverse complement of 3’-flanking region of *ALD-A*). All the three PCR products were purified and used as a template in the final PCR and amplified using a primer pair 5’- GGACTGTTCAATTTGAAGTCGATGCTGACG-3’ and 5’-GGAACTGGAGGCTTCCGCAGCAAACTCTC-3’ to obtain a 3.023 kb product consisting of zeocin expression cassette along with promoter and terminator of *ALD-A*. *GS115* strain was transformed with *Zeo*^*R*^ expression cassette and zeocin resistant colonies were selected. Deletion of *ALD-A* was confirmed by PCR using gene-specific primers.

For the deletion of *ALD-D*, 0.989 kb of promoter was amplified from *K. phaffii* genomic DNA by using primer set 1F 5’- CCAAAATGGTAACAACGTTCAAGTAAC-3’ and 1R 5’-GCTATGGTGTGTGGGGGATCCGCACAAAGTGAAGGGGAAGATAAGAGC −3’ (962 to 984 bp of *GAPDH* promoter in reverse complement, −9 to −32 bp of *ALD-D* promoter in reverse complement). *Zeo*^*R*^ expression cassette of 1.2 kb was amplified from pGAPZA vector using the primer pair 2F 5’-GCTCTTATCTT CCCCTTCACTTTGTGCGGATCCCCCACAACCATAGC-3’ (9 to −32 bp of *ALD-D* promoter and +962 to +984 bp of pGAPZA vector) and 2R 5’-GGCATATAGCAGGAGAGCTATTCCCTTTGCTCACATGTTGGTCTCCAGCTTG-3’ (+1488 bp to +1514 bp in reverse complement of 3’-flanking region of *ALD-D*, +2137 to +2159 bp in reverse complement of pGAPZA vector). 1 kb terminator sequence or 3’-flanking region of *ALD-D* promoter was amplified by using primer set 3F 5’-CAAGCTGGAGACCAACATGTGAGCAAAGGGAATAGCTCTCCTGCTATATGCC-3’ (+2137 to +2159 bp of *P*_*GAPZ-A*_ vector, and (+1488 bp to +1514 bp of region of *ALD-D* gene) and 3R 5’-TCAGAGACGATCTTCTCTTACGGG-3’ (reverse complement of +2465 to +2488 of *ALD-D* terminator region). The final products of *ALD-D* promoter, zeocin and *ALD-D* terminator was purified and used as a template to make knockout construct of *ALD-D*. The final knockout construct was amplified by using primer set 1F 5’- CCAAAATGGTAACAACGTTCAAGTAAC-3’ and 3R 5’- TCAGAGACGATCTTCTCTTA CGGG-3’. *GS115* strain was transformed with the final PCR product and antibiotic resistant colonies were screened for zeocin resistance. Deletion of *ALD-D* was confirmed by PCR by using gene-specific primers.

#### GS115-ALD-A^Myc^, Δmxr1-ALD-A^Myc^

The gene encoding ALD-A along with 1.0 kb of its promoter was cloned in-into *pIB3* vector (#25452, Addgene) and transformed into *GS115* and *Δmxr1.* The following primer pair was used: 5’- TCCC<u>CCCGGG</u>ATTGGAGAAGACAATGAAT CTG-3’ and 5’-CCG<u>CTCGAG</u>CTACAGG TCTTCTTCAGAGTCAGTTTCTGTTCCTTATGTCAGGAGTGTAAGC 3’ (*XmaI* and *XhoI* sites in the primers are underlined). The reverse primer encoded the Myc tag. The PCR product was cloned into *pIB3* vector, the recombinant plasmid was linearized using *SalI* and transformed into *GS115* and *Δmxr1*. Recombinant clones were selected by plating on YNBD His^-^ plates and clones expressing Myc-tagged ALD-A (ALD-A^Myc^) were identified by western blotting using anti-Myc antibody.

#### GS115-P_ALD-A_GFP, Δmxr1-P_ALD-A_GFP

These strains express GFP from 1.0 kb of *ALD-A* promoter and were constructed as described below: Three different PCR reactions were carried out for the construction of *P*_*ALD-A*_*GFP* expression cassette. In the first PCR reaction, 1 kb *ALD-A* promoter was amplified from genomic DNA of *GS115* strain of *K. phaffii* using a primer pair 1F (5’CG<u>GGATCCA</u>TTGGAGAAGACAATGAATCTGAC-3’, (BamHI site is underlined) and 1R (5’CTCCTTTACTAGTCAGATCTACCATGGATAAAGGTAAGGGAAAAAAGCAAGTG-3’) +1 to +25 bp of the gene encoding GFP and −1 to −28 bp of *P*_*ALD-A*_). In the second PCR reaction, a 714 bp region of *GFP* gene was amplified from pREP41GFP vector using the primer pair 2F (5’- CACTTGCTTTTTTCCCTTACCTTTATCCATGGTAGATCTGACTAGTAAAGGAG-3’, −1 to −28 bp of *P*_*ALD-A*_) and +1 to +25 bp of the gene encoding GFP) and 2R (5’- CCG<u>CTCGAG</u>CTAGTGGTGGTGGCTAGCTTTG-3’) The XhoI site is underlined. In the third PCR reaction, the PCR products from the first two reactions were used as templates and amplified using the 1F and 2R primers to get the *P*_*ALD-A*_*GFP* expression cassette, which was digested with BamHI and *XhoI* and cloned into pIB3 (#25452, Addgene) to generate *pIB3-P*_*ALD-A*_*GFP*. The generated expression vectors were linearized with *SalI* and transformed into *Komagataella phaffii GS115* and *Δmxr1* strains and plated on YNBD-His^−^ agar plates, positive colonies were screened using anti-GFP antibody.

#### GS115-ALD-A-M1^GFP^, GS115-ALD-A-M2^GFP^, GS115-ALD-A-M3^GFP^, GS115-ALD-A-M^*GFP^ and GS115-ALD-A-M2,3^GFP^

*GS115* expressing GFP from 1.0 kb *ALD-A* promoter in which each of the three MXREs (M1, M2, M3), two MXREs and all the three MXRES (M*) are mutated were generated as follows: *P*_*ALD-A*_*GFP* plasmid with MXRE-M1 was generated as follows: *ALD-A* promoter was PCR amplified using primer pair 1F (5’-CG<u>GGATCCATT</u>GGAGAAGACAATGAATCTGAC −3’, (*BamHI* site is underlined) and 1R (5’-CCCTCTCCAAAAGAGGGACGATGTAG ATGAGAACCGTTGAGCGGAATCATGGGTG-3’). In another PCR, *GFP* was amplified using primer pair 2F (5’-CACCCATGATTC CGCTCAACGGTTCTC<u>A</u>TCTACATCGTCCCTCTTTTGGAGAGGG-3’ (mutation is underlined) and 2R (5’- CCG<u>CTCGAG</u>CTAGTGGTGGTGGCTAGCTTTG-3’) (The *XhoI* site is underlined). In the third PCR reaction, the PCR products from the first two reactions were used as templates and amplified using the 1F and 2R primers. Thereafter, the amplicon was digested with *BamHI* and *XhoI* and cloned into *pIB3* (Addgene, USA) to generate a plasmid carrying mutation in MXRE1 (*pP*_*ALD-A*_*GFP-M1*). Similar strategy was used to generate the plasmids, *pP*_*ALD-A*_*GFP-M2 pP*_*ALD-A*_*GFP-M3, pP*_*ALD-A*_*-GFP-M2,3* and *pP*_*ALD-A*_*GFP-M*\* carrying mutations in MXRE2 alone, MXRE3 alone or both MXRE 2 and 3 or all three MXREs respectively using appropriate 1R mutant reverse primers. Plasmids were linearized with *SalI* and transformed into *GS115* to obtain the strains GS115-*ALD-A-M1*^*GFP*^, *GS115-ALD-A-M2*^*GFP*^, *GS115-ALD-A-M3*^*GFP*^, *GS115-ALD-A-M2,3*^*GFP*^ and *GS115-ALD-A-M*\*^*GFP*^ strains.

#### GS115-P_AOX1_GFP, Δald-a-P_AOX1_GFP

*GS115* and *Δald-a* strains expressing GFP from 1.0 kb of *AOX1* promoter were generated as follows: *AOX1* promoter (−1002 bp) was amplified by PCR using *K. phaffii* genomic DNA as a template and the primer pair 1F (5’- CGG<u>GGTACC</u>TCATGTTGGTATTGTGAAAT AGACGCAGATC-3’, −1002 to −971 bp of *P*_*AOX1*_) and 1R (5’-CTCCTTTACTAGTCAGATCTA CCATCGTTTCGAATAATTAGTTGTTTTTTGATC-3’, +1 to +25 bp of the GFP ORF and −1 to −29 bp of *P*_*AOX1*_). The *KpnI* site is underlined. In the second PCR reaction, a 714 bp region of *GFP* gene was amplified from pREP41GFP vector using the primer pair 2F (5’- GATCAAAAAACAACTAA TTATTCGAAACGATGGTAGATCTGACTAGTAAAGGAG-3’, −1 to −29 bp of *P*_*AOX1*_ and +1 to +25 bp of the *GFP*) and 2R (5’- CCG<u>CTCGAG</u>CTAGTGGTGGTG GCTAGCTTTG-3’) The *XhoI* site is underlined. In the third PCR reaction, the PCR products from the first two reactions were used as templates to amplify *P*_*AOX1*_*GFP* using 1F and 2R primers. The final PCR product was digested with *KpnI* and *XhoI* and cloned into pIB3 vector (addgene, cat No. 25452) to yield *pIB3-P*_*AOX1*_*GFP*. The recombinant plasmid was linearized with *SalI* and transformed into *GS115* and *Δald-a* strains. GFP expressing clones were identified by western blotting using anti-GFP antibodies.

## Data availability statement

All data presented and discussed are contained within the manuscript.

## Footnotes / Funding

This work was supported by the J. C. Bose Fellowship grant SB/S2/JCB-025/2015 awarded by the Science and Engineering Research Board, New Delhi, India and the research grant BT/PR30986/BRB/10/1751/2018 awarded by the Department of Biotechnology, New Delhi, India (to P.N.R). Funding from the Department of Science and Technology Fund for Improvement of S&T Infrastructure in Higher Educational Institutions (DST-FIST), the University Grants Commission and the Department of Biotechnology (DBT)-Indian Institute of Science partnership program is acknowledged. K.K.R is a recipient of junior and senior research fellowship of Indian Council Medical Research, New Delhi, India.

